# Two-step interphase microtubule disassembly aids spindle morphogenesis

**DOI:** 10.1101/126342

**Authors:** Nunu Mchedlishvili, Helen K. Matthews, Adam Corrigan, Buzz Baum

## Abstract

Entry into mitosis triggers profound changes in cell shape and cytoskeletal organisation. Here, by studying microtubule remodelling in human flat mitotic cells, we identify a two-step process of interphase microtubule disassembly. First, a microtubule stabilizing protein, Ensconsin, is inactivated in prophase as a consequence of its phosphorylation downstream of Cdk1/CyclinB. This leads to a reduction in interphase microtubule stability that may help to fuel the growth of centrosomally-nucleated microtubules. The peripheral interphase microtubules that remain are then rapidly lost as the concentration of tubulin heterodimers falls following dissolution of the nuclear compartment boundary. Finally, we show that a failure to destabilize microtubules in prophase leads to the formation of microtubule clumps, which interfere with spindle assembly. Overall, this analysis highlights the importance of the stepwise remodelling of the microtubule cytoskeleton, and the significance of permeabilization of the nuclear envelope in coordinating the changes in cellular organisation and biochemistry that accompany mitotic entry.

## Introduction

The goal of mitosis is the equal segregation of genetic material into two daughter cells. To achieve this, animal cells undergo profound changes in cell organisation. Cells round up, chromosomes condense, and the permeability of the nuclear envelope increases (a process we term NEP), leading to mixing of nucleoplasm and cytoplasm. At the same time, the array of long interphase microtubules is replaced by a population of short and highly dynamic centrosome-nucleated microtubules, which go on to form the mitotic spindle – the structure responsible for chromosome segregation.

Microtubule cytoskeleton remodelling starts before NEP with a dramatic increase in microtubule nucleation at centrosomes [1], driven by the recruitment and local activation of the gamma tubulin ring complex (gamma TuRC) [2–5]. While this burst of centrosomal microtubule nucleation is easily visible in most cell types, careful quantification of microtubule polymer levels have suggested that total cellular levels of the tubulin polymer remain relatively constant during this period, before abruptly dropping at the transition from prophase to prometaphase with NEP [6]. Once cells are in prometaphase, highly dynamic microtubules emanating from centrosomes search cell space for kinetochore attachment sites to capture chromosomes [7–9]. This stabilizes the microtubules, leading to a rise in tubulin polymer levels during mitotic spindle formation [6, 10]. Close to chromosomes, microtubule polymerisation is aided by a local Ran-GTP gradient [11–17]. The final size of the spindle is then determined by microtubule regulators [18] together with the total available pool of tubulin heterodimers [19].

The wholesale change in cell state at mitotic entry is driven by the activation of Cdk1/CyclinB complex - the master regulator of mitosis [20]. FRET-based quantitative measurements in mammalian somatic cells revealed that Cdk1/CyclinB complex first becomes activated in the cytoplasm around 20 min before NEP [21]. The activity of Cdk1/CyclinB then increases rapidly through the action of positive feedback loops [22], peaking around 10 min after the onset of prometaphase [21]. This change in Cdk1/CyclinB activity is accompanied by a change in its localization. At early stages the complex is seen at elevated levels in the cytoplasm, where it is concentrated at centrosomes in many systems [23, 24]. Then, as levels of Cdk1/CyclinB increase, it enters into the nucleus [25], where it triggers the onset of NEP [26]. Once the nuclear envelope has been permeabilized cells are committed to mitosis [27].

It is generally assumed that the differences in microtubule structure and dynamics that accompany the transition from interphase to mitosis result from phosphorylation-induced changes in the activities and/or binding capacities of stabilizing and destabilizing microtubule-associated proteins (MAPs) downstream of Cdk1/CyclinB activation [28–32]. However, given the gradual increase in Cdk1/CyclinB complex activity [21], it is hard to reconcile this view with the switch-like change in overall levels of microtubule polymer, which are reported to occur during the transition from prophase to prometaphase [6]. This suggests that there is more at play. Moreover, while mitotic spindle assembly is one of the best-studied processes in cell biology, the process by which the interphase microtubule array is remodelled at mitotic entry to give rise to the mitotic spindle remains unclear; as does its functional significance. This is in part due to the difficulties of studying microtubules in cells as they round upon entry into mitosis.

In this paper, we shed new light on this process. A detailed quantitative analysis of microtubule polymer levels in living cells reveals that interphase microtubule disassembly occurs in two sequential steps. First, interphase microtubules are gradually lost as the result of phosphorylation-induced inactivation of the microtubule stabilizer Ensconsin in prophase. This is followed by a sudden loss of tubulin polymer at prometaphse, which is triggered, at least in part, by the reduction in the concentration of tubulin heterodimers that accompanies permeabilization of the nuclear envelope. Finally, we demonstrate the importance of prophase microtubule destabilization in preventing clumps of interphase microtubules interfering with mitotic spindle assembly.

## Results

### The interphase microtubule cytoskeleton is remodelled in two discrete steps at mitotic entry

In order to explore the kinetics of microtubule remodelling upon entry into mitosis, we began by monitoring changes in the microtubule cytoskeleton organization relative to loss of the nuclear/cytoplasmic compartment boundary (NEP) in HeLa cells stably expressing histone-2B-RFP and mEGFP-α-tubulin [33]. To better visualize microtubule remodelling during this period, cells were transfected with a constitutively activated version of small GTPase Rap1 (here called Rap1*), which prevents focal adhesion disassembly [34, 35] (Figure 1 A). While Rap1* expression prevented mitotic rounding, it did not appear to significantly alter the timing of microtubule remodelling (data now shown). This analysis revealed several aspects of microtubule remodelling. First, we observed a steady accumulation of mEGFP-α-tubulin at centrosomes in prophase (Figure 1 A, B). As previously described [36], this process of centrosome maturation, leading to an increase in the capacity of centrosomes to nucleate microtubules, is driven by the local action of mitotic kinases. Centrosome maturation was accompanied by the gradual disassembly of peripherally-localized interphase microtubules (Figure 1 C). Parallel changes in cytoplasmic microtubule organisation continued up until NEP. This triggered a dramatic and sudden loss of microtubule polymers over the course of a few minutes (Figure 1 A-C).

**Figure1.**
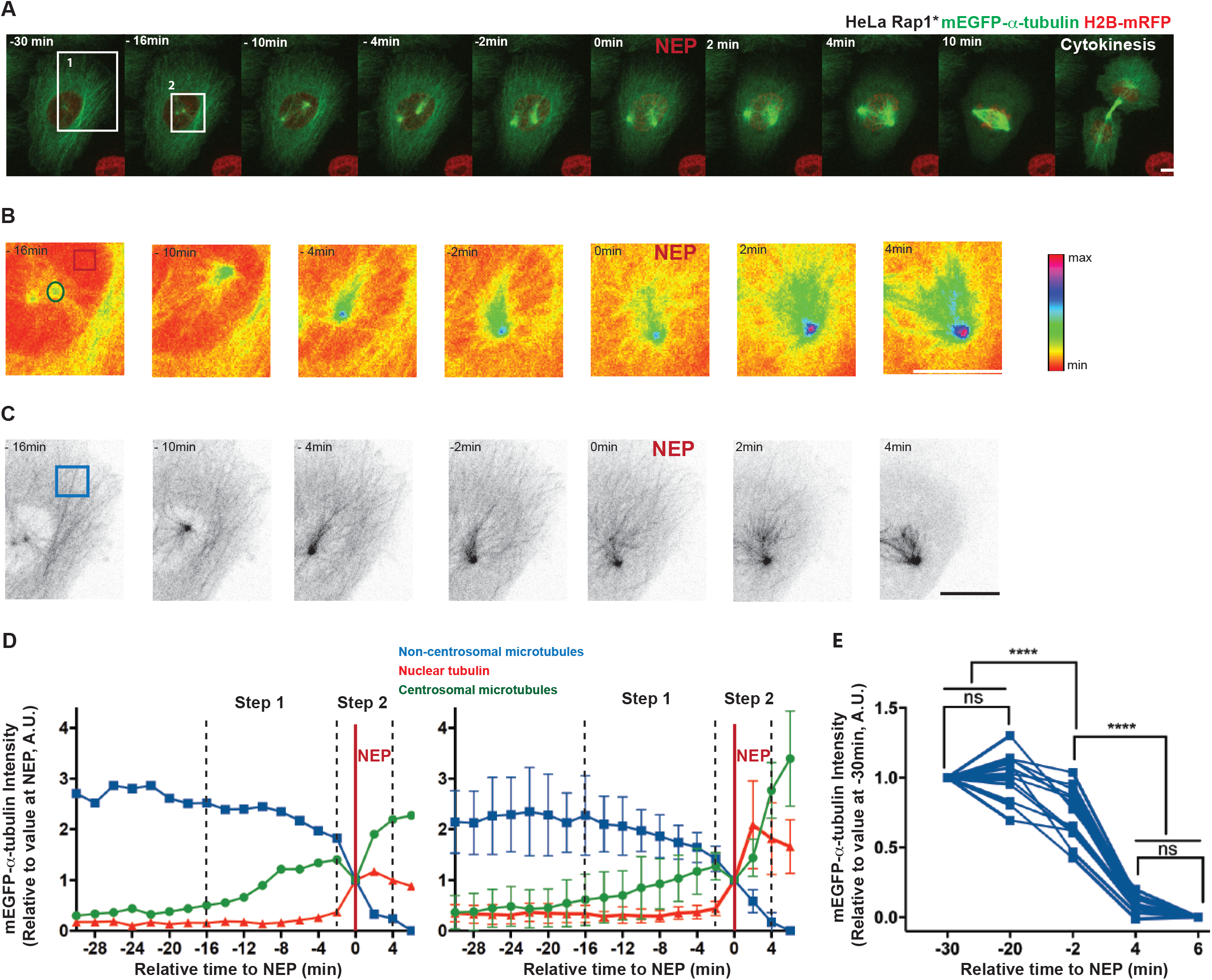
Disassembly of interphase microtubules begins prior to NEP and is accelerated at NEP. **A**) Representative time-lapse confocal images (x-y maximum projection) of a HeLa cell stably expressing H2B-mRFP (to visualize chromosomes) and mEGFP-α-tubulin (to visualize microtubules and NEP), and transiently overexpressing Rap1* (to keep cell flat as it enters mitosis). Inserts show regions zoomed in B and C. **B**) Higher magnification (sum projection of sections of mEGFP-α-tubulin around the centrosome, pseudo-color, spectra LUT) of boxed region 2 indicated in A showing how mEGFP-α-tubulin levels at the centrosome rise prior to NEP. Inserts indicate regions used for quantifications: green (centrosomal microtubules), red (nuclear tubulin). **C**) Higher magnification (maximum projection of basal sections of mEGFP-α-tubulin, inverted grayscale) of region 1 above showing that non-centrosomal microtubule disassembly is triggered before NEP and accelerates during loss of the nuclear/cytoplasmic compartment boundary. Insert indicates region used for quantifications. **D**) Changes in median centrosomal and non-centrosomal microtubule intensity relative to NEP for H2B-mRFP mEGFP-α-tubulin HeLa cell transiently overexpressing Rap1* (shown in A-C, left), and for 5 equivalent cells from 2 independent experiments (right). Median intensity of mEGFP-α-tubulin signals was calculated within a 15X15 pixel circle around the centrosome (green line, as indicated in B), a 10×10 pixel box within the nucleus (red, as indicated in B), and a 30×30 pixel box at the cell periphery (blue, as indicated in C). Time point 0 represents NEP. Graph shows means and SD. **E**) Changes in non-centrosomal microtubule levels relative to NEP. Measurements show median of mEGFP-α-tubulin signal in a 30X30 pixel box at two locations at the periphery of a cell as shown in Figure S1A at 30, 20, 2 min before NEP and 4, 6 min after NEP in H2B-mRFP mEGFP-α-tubulin HeLa stable cell line transiently overexpressing Rap1* (13 cells (include 5 cells from 1D), 4 independent experiments). Repeated Measures ANOVA, Tukey’s multiple comparisons test with a single pooled variance, **** P<0.0001. Scale bars represent 10μm.

To gain a quantitative measure of changes in microtubule polymer levels during entry into mitosis, we followed changes in the intensity of the mEGFP-α-tubulin polymer signal at centrosomes, using the rise in nuclear mEGFP-α-tubulin as a marker of NEP (Figure 1 D, Figure S1 E); and mEGFP-α-tubulin in neighbouring cells to control for bleaching (Figure S1 A-C). At the same time, we used mEGFP-α-tubulin to determine the kinetics of disassembly of interphase microtubule at the cell periphery (beyond the reach of visible microtubules emanating from the centrosome) (Figure 1 A, C, D, E). Since this population of interphase microtubules is not physically connected to the centrosome in prophase, we also refer to them as “non-centrosomal microtubules”. In line with our qualitative observations, this analysis revealed that the increase in the mEGFP-α-tubulin signal at centrosomes begins ~16 min before NEP (Figure 1 D, Figure S1 E), and is accompanied by a steady decrease in levels of mEGFP-α-tubulin polymer (a 33 ± 17% decrease in mEGFP-α-tubulin intensity over 14 min) at the cell periphery (Figure 1 D). We confirmed these findings using the local variance of mEGFP-α-tubulin intensity as an alternative method by which to quantify levels of tubulin polymer above the background monomer signal (Figure S1 D). Strikingly, these gradual changes in the levels of tubulin polymer during prophase were abruptly altered at NEP. The sudden increase in nuclear mEGFP-α-tubulin at NEP was accompanied by a transient reduction in the levels of microtubule polymer at centrosomes and the rapid loss of residual microtubules from the cell periphery (Figure 1 A-E, Figure S1 E).

Based on these results, we conclude that the microtubule cytoskeleton is reorganised in two discrete steps at mitotic entry. The first step, during prophase, is characterized by a slow partial depolymerisation of interphase microtubules at the cell periphery. This is accompanied by the nucleation of microtubules from centrosomes. Since these processes occur in parallel, total levels of tubulin polymer do not markedly change during this period. Then, coincident with the loss of the nuclear-cytoplasmic compartment barrier, there is a sudden reduction in the total levels of microtubule polymer (Figure 1 A-C). Both these measurements are in line with those of previous studies, validating our approach [6].

### Step 1: Cdk1/CyclinB dependent removal of Ensconsin from microtubules triggers non-centrosomal microtubule depolymerisation during prophase

To elucidate the molecular mechanisms that govern microtubule disassembly at the entry of mitosis, we focused first on events during prophase. In order to test whether the loss of interphase microtubules during this period is an indirect consequence of the growth of centrosomal microtubules, e.g. via competition for a common tubulin heterodimer pool, we monitored the loss of interphase microtubules in cells that fail to nucleate centrosomal microtubules as the result of RNAi-mediated silencing of Cep192 [37, 38] (Figure 2 A). Importantly, since Cep192 silencing did not significantly alter the disassembly kinetics of peripheral interphase microtubules (Figure 2 A-C, Figure S2 A), this analysis shows that the two processes, interphase microtubule loss and centrosome maturation, are regulated independently of one another.

**Figure 2.**
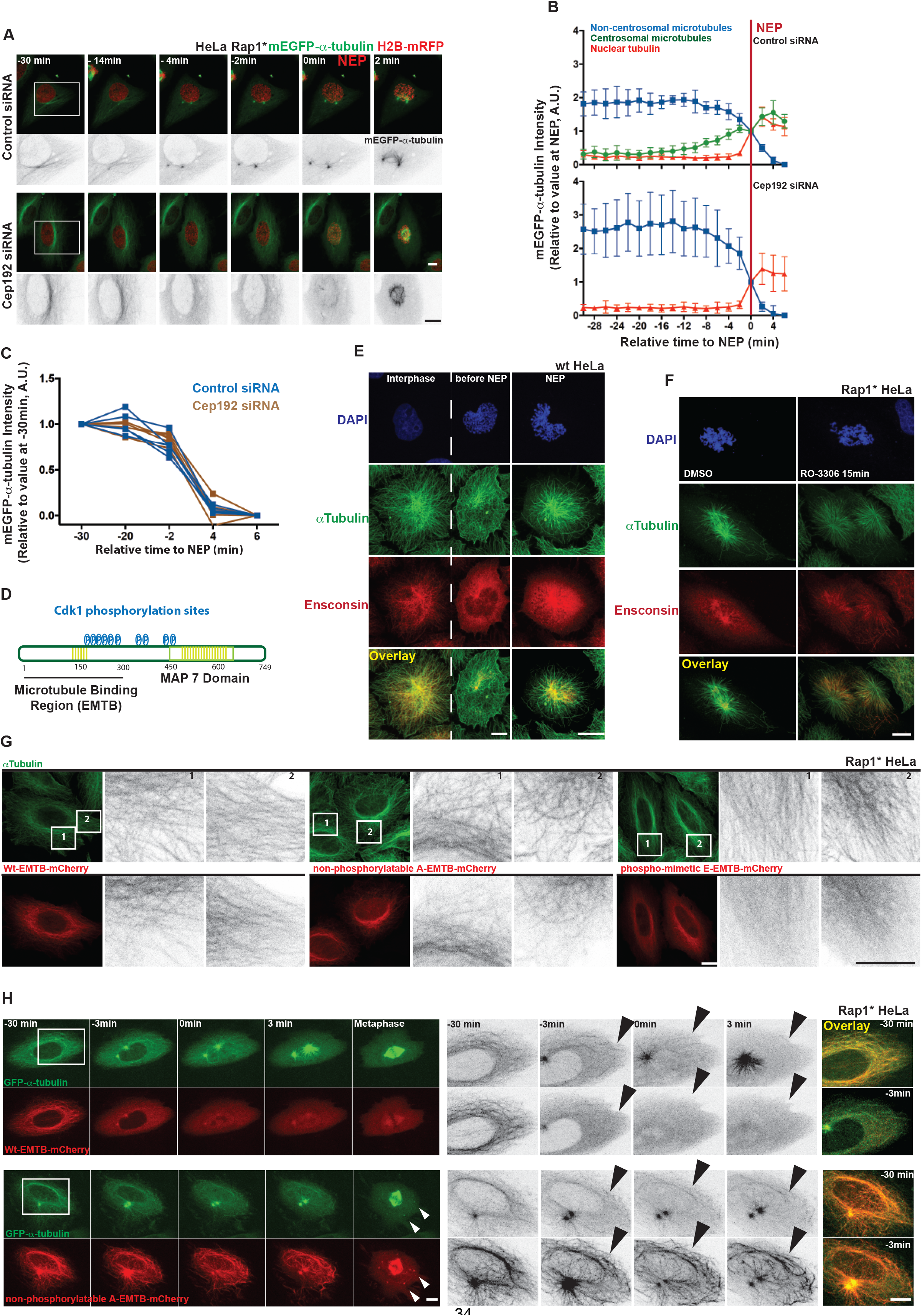
Removal of Ensconsin by Cdk1/CyclinB activation triggers non-centrosomal microtubule depolymerisation before NEP. **A**) Representative time-lapse confocal images (x-y maximum projection) of HeLa cells during mitotic entry stably expressing H2B-mRFP and mEGFP-α-tubulin, and transiently overexpressing Rap1* treated with control siRNA (upper panel) or Cep192 siRNA (lower panel). Inserts show regions zoomed in inverted grayscale. **B**) Changes in centrosomal and non-centrosomal microtubule levels relative to NEP for control siRNA (upper panel, 5 cells) and for Cep192 siRNA (lower panel, 5 cells) 2 independent experiments. Measurements were made as in Figure 1 D, except mean of α-tubulin-GFP signal around a centrosome was calculated semi-automatically in H2B-mRFP mEGFP-α-tubulin HeLa stable cell line transiently overexpressing Rap1* (see Materials and Methods for details). Graphs show means and SD. **C**) Changes in non-centrosomal microtubule levels relative to NEP. Median of α-tubulin-GFP signal measured as described in Figure 1 C for control siRNA (blue, 5 cells) and Cep192 siRNA cells (brown, 5 cells), 2 independent experiments as in 2 B. **D**) Schematic representation of Ensconsin structure. Coiled Coil domains are shown in light green. **E**) Representative confocal images (x-y maximum projection) of fixed HeLa cells stained to show that Ensconsin is removed from microtubules in prophase. Ensconsin in red, α-Tubulin in green and DAPI in blue (11 prophase cells, >30 interphase cells, 2 independent experiments). **F**) Representative confocal images of fixed HeLa cells overexpressing Rap1* stained to show that Ensconsin re-localizes at the microtubules upon Cdk1 inhibition with R0-3306. Ensconsin in red, α-Tubulin in green and DAPI in blue (9 DMSO and 10 R0-3306 cells, 2 independent experiments). **G**) Representative confocal images of fixed HeLa cells overexpressing Rap1* stained to show that overexpressed Wt-EMTB-mCherry as well as a non-phosphorylatable mutant (A-EMTB-mCherry) remain bound to microtubules in interphase, whereas the phospho-mimetic (E-EMTB-mCherry) remains largely cytoplasmic. α-Tubulin in green, mCherry in red. Inserts show regions zoomed in inverted grayscale (9 cells Wt-EMTB-mCherry, 14 cells A-EMTB-mCherry, 10 cells E-EMTB-mCherry, 1 experiment). **H**) Representative time-lapse confocal images (x-y maximum projection) of flat (Rap1*) HeLa cells stably expressing GFP-α-tubulin and wild type Ensconsin microtubule binding domanin (Wt-EMTB-mCherry, upper panel) or its non-phosphorylatable mutant (A-EMTB-mCherry, lower panel) to show that the non-phosphorylatable mutant stays associated with microtubules during prophase, leading to delay in microtubule disassembly at mitotic entry. Inserts show regions zoomed in inverted grayscale. White arrows indicate to microtubule clumps formed due to a failure to remove Ensconsin from microtubules before NEP. Black arrows indicate to interphase microtubules just before or after NEP (18 cells Wt-EMTB-mcherry, 15 cells A-EMTB-mcherry, 5 independentexperiments). In overlay images, signal intensities were adjusted to remove cytoplasmic background signal. Scale bars represent 10μm.

Cdk1/CyclinB complex activity rises at around the time that interphase microtubules begin to depolymerize [26]. Since Cdk1/CyclinB is thought to mediate many of the structural changes that accompany entry into the mitosis, including microtubule cytoskeleton remodelling [28–32]), in our search for regulators of this loss of interphase microtubules at the entry into mitosis, we focused our attention on conserved regulators of microtubules that are potential substrates of Cdk1/CyclinB. This led us to explore the function of Ensconsin in this system. This choice was based upon the fact that Ensconsin (also called MAP-7 and E-MAP-115) (Figure 2 D) is a microtubule-associated protein that is phosphorylated in mitosis [39]. Moreover, while both fly and human Ensconsin homologues function to stabilize microtubules [40] [41] [42], the hyper-phosphorylated, mitotic form of the protein is unable to bind to microtubules *in vitro* and in HeLa cells [39]. Further, a quantitative proteomics study identified Ensconsin as a protein that preferentially binds interphase microtubules [43]. Although data from some cells types does not entirely fit this model [44], overall this previous work suggested Ensconsin as a potential regulator of changes in microtubule dynamics upon passage from interphase into mitosis.

To begin our analysis of the localization, regulation and function of Ensconsin, we used immunofluorescence to visualize changes in the localization of the endogenous protein during mitotic entry (Figure 2 E). The specificity of the antibody was confirmed via western blot using extracts prepared from cells treated with three different siRNAs targeting Ensconsin mRNA and by immunofluorescence (Figure S2 B-C). As previously reported [39], Ensconsin decorated interphase microtubules in untreated HeLa cells, but was lost from microtubules in early prophase (Figure 2 E). The same result was obtained in MCF10A cells (Figure S2 D). Since the Ensconsin microtubule binding domain (EMTB) was previously shown to have the same subcellular localization as the full-length protein [45], and to be sufficient to stabilize microtubules against drug-induced disassembly *in vivo* [40], we used this shorter version of the protein to dissect the mechanisms controlling the change in its localization.

Like the endogenous protein, when EMTB-mCherry was expressed in flat Rap1* HeLa cells stably expressing GFP-α-tubulin [46], it localized to microtubules in interphase (Fig 2 H, top panel), but was lost from microtubules as cells entered mitosis (Figure 2 H, top panel) - a result that was confirmed in fixed cells (Fig 2 G, Figure S2 E, top panel). Importantly, the dissociation of EMTB from interphase microtubules was found to precede their depolymerisation (Figure 2 H, top panel, Figure S2 E, top panel).

To determine if the removal of Ensconsin from microtubules at the onset of mitosis is mediated via phosphorylation downstream of the mitotic kinase Cdk1/CyclinB (Figure 2 D), we then treated HeLa cells overexpressing Rap1* with the Cdk1 inhibitor RO-3306 (or DMSO) for 15 minutes (Figure 2 F). Cells were fixed and stained for α-tubulin, Ensconsin and DNA. As expected, in cells treated with the Cdk1 inhibitor that were forced to exit mitosis, as indicated by the de-condensation of chromatin and the increase in tubulin polymer levels, Ensconsin was found associated with microtubules (Figure 2 F). In light of this, to determine whether the Cdk1/CyclinB-dependent dissociation of Ensconsin from microtubules is mediated by phosphorylation, we manually searched the protein sequence for mitotic kinase consensus sites (Figure 2 D). This identified ten Cdk1 sites (T/SP) in the protein, six of which were present in the microtubule-binding domain of Ensconsin, which we showed mimics the behaviour of the full-length protein. Importantly, all have been previously identified as sites of phosphorylation in mitotic cells using mass spectrometry [47, 48] (just one of these sites was found to be phosphorylated in both mitosis and G1[47]). In addition, we identified four potential Nek2 sites in the region. To determine their function, we mutated these ten putative mitotic kinase sites in the context of EMTB-mCherry to generate non-phosphorylatable (here called A-EMTB-mCherry) and phospho-mimetic variants (here called, E-EMTB-mCherry). Constructs encoding wildtype, A-mutant and E-mutant versions of EMTB-mCherry were transiently expressed in HeLa cells. Cells were then fixed and labelled with anti-α-tubulin, anti-mCherry antibodies. Whereas both the wildtype and the A-EMTB-mCherry constructs decorated interphase microtubules, the E-EMTB protein remained diffuse in the cytoplasm (Figure 2 G). Furthermore, while wildtype EMTB-mCherry was lost from microtubules in prophase, during this period the A-mutant EMTB-mCherry remained tightly associated with microtubules (Figure S2 E). These data support the hypothesis that, upon entry into mitosis, Ensconsin’s association with microtubules is regulated by phosphorylation within the microtubule binding domain.

Next to test whether this phospho-regulation has an impact on interphase microtubule disassembly we transfected a HeLa GFP-α-tubulin stable cell line expressing Rap1* with either a wildtype or an A-mutant version of the EMTB-mCherry construct. We then monitored the remodelling of the microtubule cytoskeletal as cells entered mitosis live. Strikingly, the continued association of mutant A-EMTB-mCherry protein with interphase microtubules was sufficient to increase their stability; so that many now persisted into prometaphase (Figure 2 H). Thus, the phosphorylation of Ensconsin within the microtubule binding domain is important for the timely disassembly of interphase microtubules during entry into mitosis.

### The function of interphase microtubule disassembly prior to loss of the nuclear/cytoplasmic compartment boundary

In many of the cells expressing A-EMTB-mCherry, the interphase microtubules that fail to become destabilized upon entry into mitosis formed clumps in prometaphase, which persisted on into metaphase. These microtubule clusters either became incorporated into the developing spindle or were maintained intact outside the spindle until the end of division (Figure 2 H, Movie S1). Interestingly, similar microtubule clumps have been previously described in mitotic cells treated with the microtubule stabilizing drug, Taxol [49–51]. To confirm these findings and to test whether microtubule clumps seen under these conditions have a similar aetiology to those seen in cells expressing A-EMTB-mCherry, we treated cells with 2nM Taxol, a dose previously shown to stabilize microtubules [52]. This was sufficient to prevent the interphase microtubule disassembly during prophase (Figure 3 A-B), leading to the formation of stable microtubule clumps, many of which failed to resolve prior to spindle formation. In some instances, this led to the formation of multipolar spindles, which underwent multipolar divisions in both flat (Figure 3 A, C) and round cells (data not shown as previously shown [51]). Interestingly, lower doses of Taxol (1nM) induced a partial stabilization of microtubules, more similar to that seen following the expression of A-EMTB-mCherry (Figure 3 A-B). This led to the stabilization of interphase microtubules, the formation of microtubule clumps during prometaphase, and to the transient formation of multipolar spindles (Figure 3 A-C) – most of which resolved prior to anaphase. These data make clear the importance of removing the population of interphase microtubules prior to assembly of the spindle.

**Figure 3.**
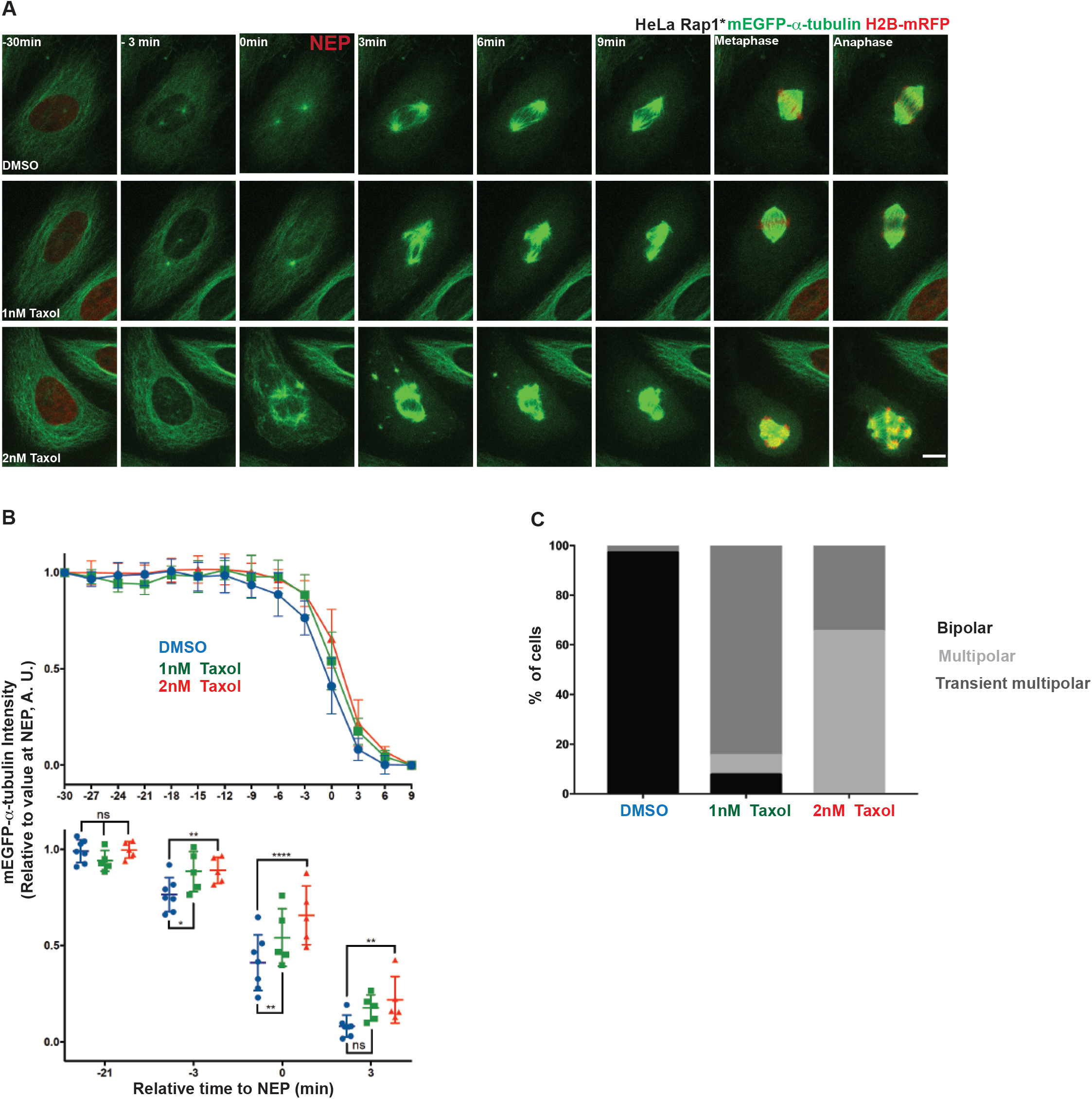
Destabilization of non-centrosomal microtubules prior to NEP is important for normal spindle assembly. **A**) Representative time-lapse confocal images (x-y maximum projection) of a flat (Rap1*) HeLa cell during mitotic progression stably expressing mEGFP-α-tubulin and H2B-mRFP (shown only at −30, during metaphase and anaphase, to better visualize microtubules), treated with DMSO (upper panel), 1nM Taxol (middle panel) or 2nM Taxol (lower panel). **B**) Changes in non-centrosomal microtubule levels relative to NEP for cells treated with DMSO (blue, 7 cells pooled 2 independent experiments), 2nM Taxol (red, 5 cells, 1 independent experiments) or 1nM Taxol (green, 5 cells, 1 independent experiments). Measurements are done as described in Figure 1 E. Graphs show means and SD. Lower panel: comparison between DMSO, 1nM and 2nM Taxol at −21, −3, 0 and 3 min relative to NEP. Repeated Measures two-way ANOVA, Dunnett’s multiple comparisons test, **** p≤0.0001, *** P≤0.001, ** P≤0.01, * P≤0.05. **C**) Quantification of % mitotic spindle defects in cells (as above) treated with DMSO (34 cells, 4 independent experiments), 1nM (13 cells, 2 independent experiments) or 2nM Taxol (26 cells, 2 independent experiments). Scale bar represents 10μm.

Having established a role for Ensconsin in the destabilization of interphase microtubules during prophase, we next wanted to determine the cause of the sudden depolymerization of remaining microtubules observed during loss of the nuclear-cytoplasmic compartment boundary (1 A-E, Figure S1 E). To improve our ability to track microtubules under these conditions, we again imaged flat HeLa cells expressing histone-2B-RFP and mEGFP-α-tubulin at a high temporal resolution (taking a frame every 0.2 min). We also pre-treated cells with STLC (to block centrosomes separation) to facilitate our ability to image both centrosomally-nucleated microtubules and peripheral interphase microtubules (Figure 4 A). Under these conditions, the marked influx of mEGFP-α-tubulin into the nucleus at NEP was precisely correlated with the loss of residual interphase microtubules and with a transient dip in the levels of mEGFP-α-tubulin polymer at centrosomes (Figure 1, Figure 4 A-B). Strikingly, microtubule polymer levels at centrosomes then quickly recovered in the ensuing minutes. This momentary reversal in the steady accumulation of centrosomally-nucleated microtubules that make up the mitotic spindle is hard to explain as a simple consequence of changes in local or global mitotic kinase activity. This led us to consider a simpler hypothesis. Since NEP is associated with the sudden dilution of cytoplasmic proteins, including the pool of tubulin heterodimers, this suggested a model whereby the sudden change in tubulin heterodimer concentration at NEP causes a relatively rapid change in the kinetics of tubulin polymer assembly. This is due to the fact that the concentration of tubulin heterodimer is a key factor in determining the dynamic behaviour of microtubules. This is true even in the case of microtubules assembled from pure tubulin *in-vitro*. In these experiments, a decrease in tubulin heterodimer concentration leads to an increase in the frequency of catastrophe events [53] and to a decrease in the rate of microtubule nucleation [54]. To determine whether the changes in tubulin heterodimer concentration that occur during NEP are of the right order to explain the observed changes in microtubule assembly, we used a nuclear-localized protein, MS2-mCherry-NLS to measure the volume of the nucleus relative to that of the entire cell during the transition into mitosis [55]. On average, NEP led to a 3-fold (±0.16) increase in the volume occupied by the MS2-mCherry-NLS signal (Figure 4 C-D), and a similar 4-fold (±1.6) decrease in MS2-mCherry-NLS concentration (Figure 4 E). Next, to measure the extent of tubulin heterodimer dilution, we performed the converse analysis in nocodazole-treated HeLa cells expressing mEGFP-labelled tubulin and histone-2B-RFP mEGFP-α-tubulin. To facilitate the accurate measurement of nuclear and cell volume under these conditions, we imaged rounded cells held in non-adhesive chambers (Figure 4 F). In these cells, NEP was associated with a decrease in the peripheral mEGFP-α-tubulin signal by an average of 18±6% (Figure 5 F-H). Taken together, these data confirm that loss of the nuclear-cytoplasmic compartment boundary is accompanied by a significant reduction in the concentration of tubulin heterodimers.

**Figure 4.**
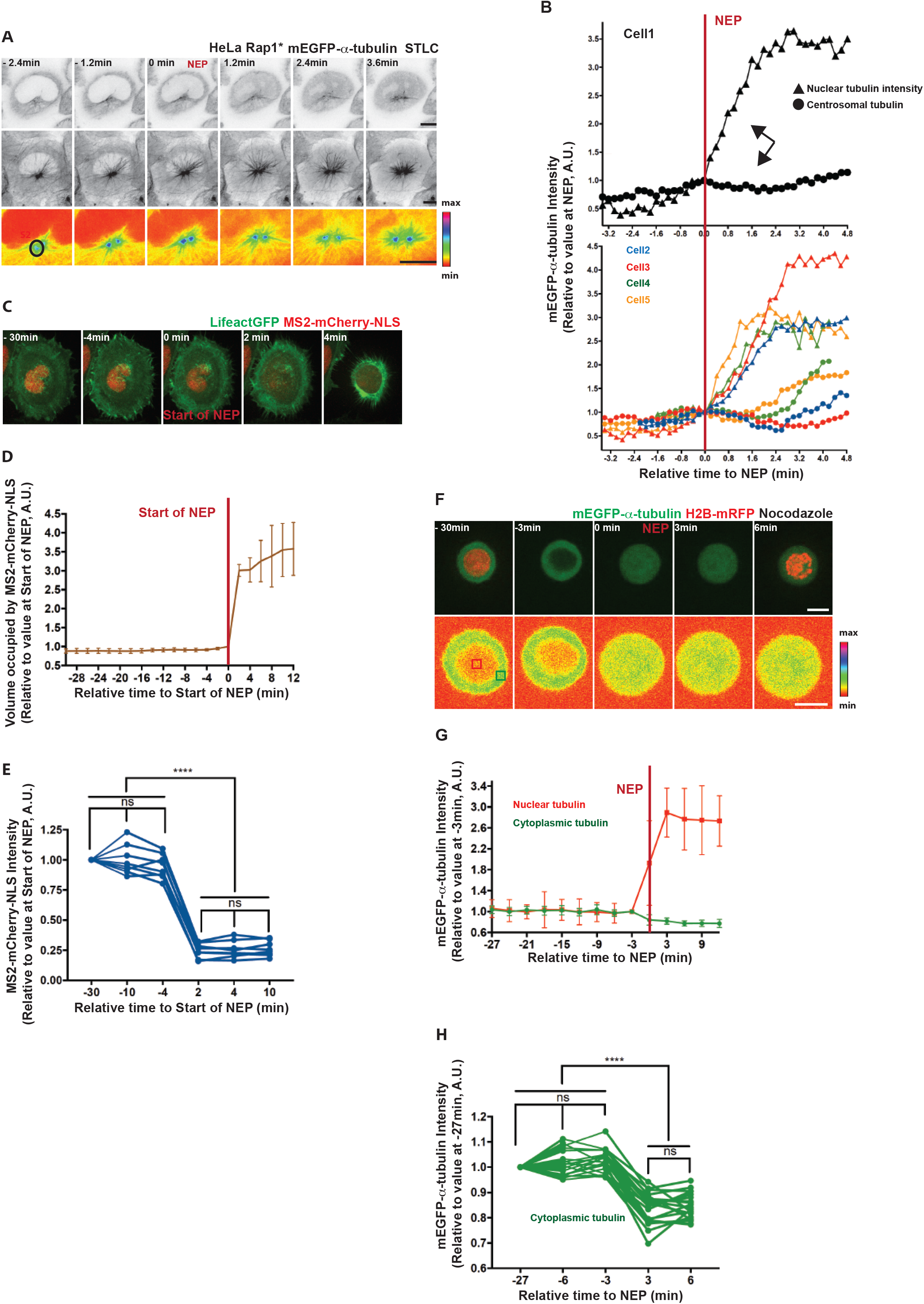
Loss of the nuclear/cytoplasmic compartment barrier is associated with the loss of tubulin polymer. **A**) Representative time-lapse confocal images of a HeLa cell during mitotic entry stably expressing H2B-mRFP (was not imaged) and mEGFP-α-tubulin, and transiently overexpressing Rap1* treated with STLC. Upper panel: upper single section (inverted grayscale) of mEGFP-α-tubulin to visualize NEP, middle panel: x-y maximum projection images of mEGFP-α-tubulin (Inverted grayscale) to visualize microtubules, lower panel: sum projection of sections of mEGFP-α-tubulin around the centrosome (pseudo-color, spectra LUT) to visualize centrosomal microtubules showing the NEP is correlated with transient decrease in mEGFP-α-tubulin intensity at centrosomes. **B**) Changes in centrosomal microtubule levels (circles) relative to NEP (nuclear tubulin, triangles) for one cell (shown in A, upper panel) and for equivalent 4 cells from 2 independent experiments (lower panel) measured similar to that described in Figure 1 D (see Materials and Methods). **C**) Representative time-lapse confocal images of a HeLa cell expressing MS2-mCherry-NLS (nuclear marker) and Lifeact-GFP during mitotic entry. **D**) Quantification of volume occupied by MS2-mCherry-NLS signal in HeLa cells similar to 4 C (8 cells, 2 independent experiments) entering mitosis. Graph shows means and SD. **E**) Quantification of MS2-mCherry-NLS intensity. Median of MS2-mCherry-NLS signal is shown for 20X20 pixel box at two locations in the nucleus per cell at 30,10, 4 min before NEP and 2, 4,10 min after NEP (8 cells, 2 independent experiments, the same cells as in 4 E). Repeated Measures ANOVA with the Greenhouse-Geisser correction, Tukey’s multiple comparisons test with individual variances computed for each comparison, **** P<0.0001. **F**) Representative time-lapse confocal images of a HeLa cell stably expressing H2B-mRFP (shown in the upper panel at -30 min and 6 min) and mEGFP-α-tubulin during mitotic entry, following treatment with Nocodazole in non-adherent chambers (upper panel: x-y maximum projection: lower panel single section, pseudocolor, spectra LUT) to show cytoplasmic tubulin dilution at NEP. **G**) Quantification of median mEGFP-α-tubulin signal in 6X6 pixel box in nucleus and at two locations at the periphery of cells as in 4F (8 cells, 1 independent experiment) entering mitosis in the non-adherent chambers filmed at high resolution using spinning disc confocal microscope. **H**) Quantification of median mEGFP-α-tubulin signal in 4X4 pixel box at four locations at the periphery of the cells similar to 4 F (20 cells, 2 independent experiments) at 27, 6, 3 min before NEP and 3, 6 min after the NEP in non-adherent chambers filmed at low resolution with wide-field microscope. Repeated Measures ANOVA with the Greenhouse-Geisser correction, Tukey’s multiple comparisons test, with individual variances computed for each comparison, **** P<0.0001. Scale bars represent 10μm.

**Figure 5.**
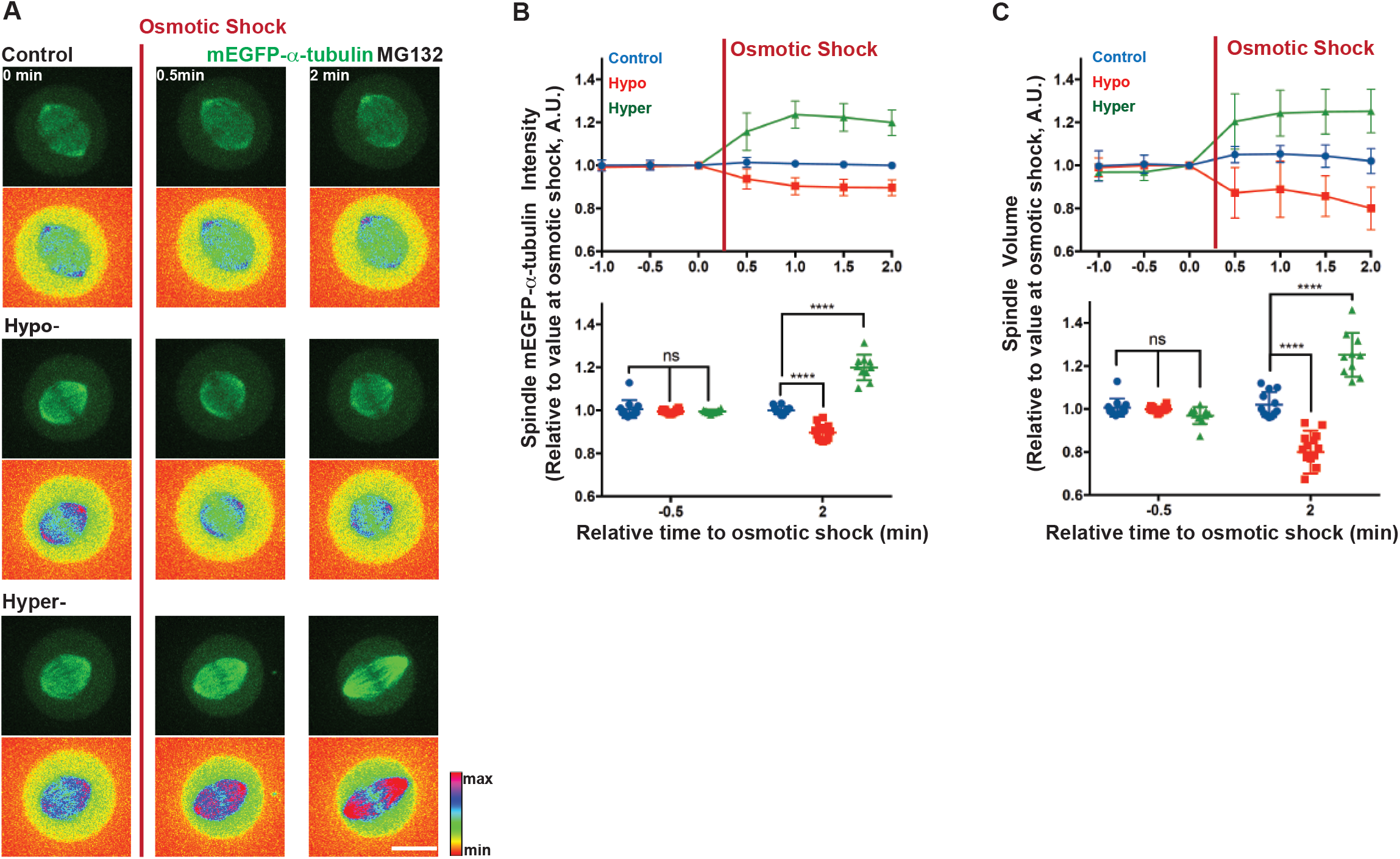
Changes in volume are sufficient to induce changes in tubulin polymer levels. **A**) Representative time-lapse confocal images (x-y maximum projection, lower panel: pseudo-color, spectra LUT) of HeLa cells stably expressing H2B-mRFP (not imaged) and mEGFP-α-tubulin showing changes in spindle mEGFP-α-tubulin intensity and spindle volume upon hypo- or hyper-osmotic shock relative to a control. Scale bar represents 10μm. **B**) Quantification of changes in spindle mEGFP-α-tubulin intensity induced by osmotic shock. Mean intensity of spindle mEGFP-α-tubulin intensity was measured by rendering mitotic spindles in 3D using Imaris software before and after control (blue, 12 cells), hypo (red, 14 cells) or hyper (green, 10 cells) osmotic shock treatment (2 independent experiments). Lower panel: comparison between −0.5 min and 2 min relative to osmotic shock treatment. Repeated Measures two-way ANOVA, Dunnett’s multiple comparisons test, **** P=0.0001. **C**) Quantification of changes in spindle volume induced by osmotic shock. Spindle volume was measured by rendering mEGFP-α-tubulin signal in 3D in Imaris before and after control (blue, 12cells), hypo (red, 14) or hyper (green, 10 cells) osmotic shock treatment (2 independent experiments). Lower panel: comparison between −0.5 min and 2 min relative to osmotic shock treatment. Repeated Measures two-way ANOVA, Dunnett’s multiple comparisons test, **** P=0.0001.

To determine how a relatively sudden 15–20% dilution of the tubulin pool is likely to affect microtubule polymer levels in mitosis, we established an assay enabling us to induce a similar, rapid reduction in the *in vivo* concentration of tubulin heterodimers using hypo-osmotic shock (Figure S3 A). While the control treatment did not trigger any significant changes in either mEGFP-α-tubulin intensity or in the cell dimeter, hypo-osmotic shock induced changes in tubulin heterodimer concentration that were quantitatively similar to those observed at NEP: a 6.3±2.5% increase in the cell diameter and a concomitant 16.2±2.8% decrease in the mEGFP-α-tubulin signal within two minutes of the shock (Figure S3 A-C). To test whether this is sufficient to reduce microtubule polymer levels, as expected under this model, we repeated this analysis in cells that had been arrested in mitosis using MG132 treatment. Under these conditions, a dilution of tubulin dimers equivalent to that observed at NEP triggered a dramatic reduction in the levels of tubulin polymer. Within two minutes of the shock, we observed a 11.4±3.7% decrease in the mEGFP-α-tubulin signal within the mitotic spindle and an accompanying 20±10% decrease in the mitotic spindle volume (Figure 5 A-C).

For the converse experiment we established a protocol to increase the tubulin dimer concentration using hyper-osmotic shock (Figure S3 A-C, see Materials and Methods). This increased the cytoplasmic mEGFP-α-tubulin signal on average by 53.7±3.6% (Figure S3 A-C) and resulted in an increase in the spindle mEGFP-α-tubulin signal by an average of 20.0±5.9%, together with a 25.2±10.2% increase in the mitotic spindle volume within two minutes of the shock (Figure 5 A-C). As these data make clear, the loss of the nuclear-cytoplasmic compartment boundary induces rapid changes in the concentration of cytoplasm that are likely to have a significant impact on a wide range of cellular processes, including microtubule disassembly. Moreover, this will be compounded by the osmotic swelling associated with NEP, which is reported to be of the order of 15–20% [56, 57].

## Discussion

### The dynamics of interphase microtubule disassembly

In this study, we explore how interphase microtubule disassembly is influenced by mitotic entry. While this is a central part of the spindle assembly process, the molecular mechanisms involved remain poorly understood because of the difficulties of imaging microtubules in cells undergo mitotic rounding. For this reason, the remodelling of the microtubule cytoskeleton at mitotic entry has been studied in most detail in Ptk1 and LLCPk1 cells [6, 58], which have few chromosomes and remain relatively flat during mitosis, rather than in human cells. Here, in order to visualize the dynamics of microtubule remodelling in live human cells, we used Rap1* overexpression to flatten HeLa cells carrying fluorescently tagged proteins. In this way we were able to analyse the dynamics of microtubule remodelling upon entry into mitosis in detail to reveal the following:

i) the activation of centrosomal nucleation is accompanied by the destabilization and gradual loss of interphase microtubules.
ii) the activation of centrosomal nucleation is not mechanistically coupled to the disassembly of interphase microtubules.
iii) the loss of the nuclear-cytoplasmic compartment boundary is associated with dilution of the tubulin heterodimer pool, with the sudden loss of residual interphase microtubules [6, 58], and with a transient dip in the levels of centrosomally-nucleated microtubules.

### The mechanism contributing to interphase microtubule disassembly

Our analysis points to there being two distinct processes at play. First, the rise of Cdk1/CyclinB activity that drives entry into mitosis induces the phosphorylation of a microtubule stabilizing protein, Ensconsin. This reduces its affinity for microtubules [39, 43], making interphase microtubules susceptible to subsequent disassembly. Although these data suggest a specific role for Cdk1/CyclinB in this regulation, the experiments performed do not exclude a role for other mitotic kinases (including Nek2). In addition, even though our data point to the importance of the timely removal of Ensconsin from interphase microtubules, it is likely that the phosphorylation of other MAPs by mitotic kinases also contributes to this process. Indeed, previous work has suggested that the phosphorylation MAP4 by Cdk1 reduces its ability to stabilize microtubules [59, 60]. Moreover, Plk1, another major mitotic kinase, has been shown to stimulate microtubule polymerization activity of MCAK, a key regulator of microtubule dynamics [61, 62]. Thus, the activation of CDK1/CyclinB is likely to trigger the remodelling of the microtubule cytoskeleton through several parallel processes. While this is the case, our data show that while the rise in mitotic kinase activity sets the stage for the depolymerisation of microtubules following the loss of nuclear/cytoplasmic compartment boundary, it is not in itself sufficient for the complete disassembly of interphase microtubules.

As our analysis makes clear, the profound changes in cytoskeletal organisation accompany loss of the nuclear-cytoplasmic compartment barrier - the event previously suggested to commit cells to mitosis [27]. This triggers a relatively rapid but transient loss of microtubule polymer from centrosomes, together with loss of the residual interphase microtubules. The transient loss of centrosomal microtubules is particularly surprising here, since this is the population of microtubules required to form the nascent spindle (Figure 1). In looking for alternative mechanisms to explain this behaviour, we focused on the role of loss of the nuclear compartment barrier itself, since this is expected to be accompanied by a relatively sudden reduction in the concentration of tubulin heterodimers [19] [53] [54] (which we measured as ~15–20%). In line with this idea, osmotic shocks designed to mirror or reverse the observed changes in tubulin concentration that accompany NEP were found to have a profound impact on microtubule polymer levels in mitotic cells. In general, the extent of the dilution in tubulin heterodimer (and other exclusively cytoplasmic proteins) accompanying mitotic entry will depend on two factors: i) the relative size of the nuclear and cytoplasmic compartments, which varies across mammalian cell types (being very large in embryonic stem cells), and ii) the extent of mitotic cell swelling, which accompanies loss of the nuclear-cytoplasmic compartment barrier [56, 57]. When combined, our data suggest that these two factors will lead to a profound and functionally significant change in tubulin concentration. In this, NEP will act in concert with alterations in the activity of microtubule associated proteins that are triggered by mitotic entry to change microtubule dynamics [18]; especially since many of these are preferentially sequestered within a single compartment (the cytoplasm or nucleus) in interphase.

### Importance of interphase microtubule disassembly

In the course of our analysis, we also tested the functional importance of this being a two-step process of microtubule destabilization. We did this by exploring the consequences of blocking the ability of mitotic kinases to induce the release of Ensconsin from microtubules in prophase. In line with this being an important event, the expression of a non-phosphorylatable form of the Ensconsin microtubule binding domain interfered with the normal process of interphase microtubule disassembly. Strikingly, the microtubules that remained formed clusters (perhaps as a result of dynein/dynactin dependent trafficking [58]), which resembled those induced by the microtubule-stabilizing compound Taxol, a drug widely used in cancer treatment [49]. These were not incorporated into the nascent spindle. In fact, under some conditions, they perturbed spindle assembly, leading to the formation of multipolar spindles and divisions – like those seen in response to clinically relevant concentrations of Taxol [63, 64]. Interestingly, in this light, these data also provide an explanation for previously published observations that Taxol only induces multipolar spindle formation when added before cells enter mitosis or during early prophase [49–51]; not when added after microtubule cytoskeleton remodelling [65, 66]. Thus, the destabilization of microtubules prior to NEP is a critical part of the process of interphase microtubule disassembly, and is a necessary prelude to spindle morphogenesis.

This two-step regulation of microtubule dynamics at mitotic entry has an additional consequence. As interphase microtubules are gradually lost upon during the course of prophase, the tubulin that is released is incorporated into microtubules nucleated during the process of centrosome maturation. These grow to long lengths, preserving the overall levels of tubulin polymer (Figure 1, [6]). While the consequences of this are not completely clear, this may aid the separation and positioning of centrosomes, since cells at this stage have yet to complete mitotic rounding [67]. Then, as a consequence of NEP, the population of long centrosomally-nucleated microtubules is quickly replaced by short, dynamic centrosomal microtubules that are used to search out and capture kinetochores. At the same time, short astral microtubules function to refine spindle positioning in the confines of the rounded metaphase cell. Thus, while it remains to be tested, it is possible that this type of two-step regulation enables centrosomal microtubules to perform distinct functions during prophase and prometaphase. The timing of nuclear envelope permeabilization may therefore be important in mediating this transition between modes of microtubule remodelling.

### Conclusion

In summary, our analysis reveals a two-step process by which interphase microtubules are disassembled upon entry into mitosis. The disassembly of interphase microtubules frees up tubulin heterodimers for its incorporation into the spindle, and aids mitosis by preventing non-centrosomal microtubules from interfering with spindle morphogenesis. This need to disassemble interphase microtubules in prophase may, in part, explain the toxicity of Taxol in dividing cancer cells [64]. Further, this study shows that the changes in cell organisation that accompany mitosis are driven both by changes in the regulation of individual proteins by phosphorylation, and by the profound changes in the make-up of the cytoplasm that accompany the transition into mitosis [68].

## Materials and Methods

### Cell Culture, RNAi, DNA Transfection, Mutagenesis, Immunoblotting, and Drug Treatments

Unlabelled HeLa Kyoto cells and HeLa stable cell lines stably expressing histone-2B-RFP and mEGFP-α-tubulin [33], and GFP-α-tubulin [46] were cultured at 37°C in a humidified incubator under 5% CO2 in Dulbecco’s Modified Eagle Medium (DMEM, Gibco) with 10% fetal bovine serum (Gibco) and 1% Pen-Strep (Sigma-Aldrich). Unlabelled MCF10A cells were cultured at 37°C in a humidified incubator under 5% CO2 in Dulbecco’s Modified Eagle Medium: Nutrient Mixture F-12 (DMEM/F12, Gibco) with 5% horse serum and 1% Pen-Strep (Sigma-Aldrich), and with the following supplements: 20ng/ml human Epithelial Growth Factor (hEGF, Roche), 0.5 mg/ml Hydrocortisone (Sigma), 100 ng/ml Cholera Toxin (Sigma), 10μg/ml Insulin (Sigma). The cell were tested and are mycoplasma free.

Lipofectamine 2000 (Invitrogen) was used for siRNA transfections according to the manufacturer’s protocol. Cells were analysed 48 hrs after transfection. siRNAs were used to knock down Ensconsin (CUACAAAGCUGCACACUCU, UCAGAGAAACGGUGAUAUA, CCAUGAAUCUUUCGAAAUA, all Dharmacon), Cep192 [38] (AGC AGC UAU UGU UUA UGU UGA AAA U (Eurofins custom designed, transfected cells were identified by abolished mEGFP-α-tubulin signal at centrosomes), and compared to a non-targeting control (AllStars Negative Control siRNA, Qiagen).

HeLa cells were transfected with pRK5-Rap1[Q63E] (Rap1*, cells transfected with Rap1* were identified by their failure to round up at the mitotic entry) [34], EMTB-mCherry (a kind gift from J. Pines lab), A/E-EMTB-mCherry (synthetized by Eurofins), where following amino acids were mutated to Alanine/Glutamic acid compared to wt-EMTB: Cdk1 sites: S169, S209, S219, T231, S254, T277; Nek2 sites: S165, S188, S202, S240 (positions refer to Ensconsin canonical sequence, UniProt identifier: Q14244-1), MS2-mCherry-NLS [55] together with LifeAct-GFP (a kind gift from Ewa Paluch’s lab) or EMTB-3XGFP (was a gift from William Bement, Addgene plasmid *#* 26741) using Fugene HD (Promega) according to the manufacturer’s instructions. Cells were analysed 24 hrs after transfection.

Depletion of Ensconsin was verified by Immunoblotting. siRNA treated cells were lysed with 1XSB (Invitrogen) were loaded onto an SDS-PAGE gel before transfer onto an Immobilon-P (Millipore) membrane by wet western blotting. Membranes were blocked in 5% BSA in TBST for 1 hr, incubated overnight at 4C with primary antibodies, and for an hour at room temperature with secondary antibodies. Antibodies were used at the following dilutions: Ensconsin (MAP7) 1:2500 (Proteintech, 13446-1-AP) and α-tubulin 1:5000 (DM1A, Sigma) and HRP-conjugated secondary antibodies 1:5000 (DAKO). Results were visualized using an ImageQuant LAS4000 system.

Drugs were used at the following concentrations: 100 ng/ml nocodozole (Sigma), 1 or 2nM Taxol (Sigma), 1μM MG132 (sigma), 10μM S-trityl-L-cysteine (STLC; Sigma), 9 μM RO-3306. Cells were incubated at least 1 hr with the drugs (except RO-3306, which was left on cells only for 15min), before live cell imaging.

### Live cell imaging

For confocal live-cell imaging, cells were seeded on glass-bottomed dishes (MatTek) coated with 10 mg/ml fibronectin (Sigma) and imaged in Leibovitz’s L-15 medium (Gibco) using an UltraView Vox (Perkin Elmer) spinning disc confocal microscope with 60X (NA 1.4) or 100X (NA 1.4) oil objective equipped with temperature controlling environmental chamber. Images were acquired using a Hamamatsu ImagEM camera and Volocity software (Perkin Elmer). Wide-field live-cell imaging was done using a Zeiss Axiovert 200M microscope with a 20Xobjective (NA 0.4) equipped with temperature controlling environmental chamber, and images acquired using a Hamamatsu Orca AG camera and Volocity software (Perkin Elmer).

For the tubulin dilution calculations at NEP, cells were seeded in PDMS chambers incubated overnight with 0.1mg/ml PEG (Poly(ethylene glycol), Sigma) dissolved in 10mM Hepes pH 7.4.

Osmotic shock was induced either by diluting imaging media by 50% with deionized water (hypo-osmotic shock) or by adding 4mM Sorbitol (Sigma) solution to the end concertation of 1.3mM (hyper-osmotic shock). For the control experiments imaging media was added.

### Immunofluorescence

For immunostaining, cells were plated on fibronectin-coated glass chambers and either fixed with 4% formaldehyde for 20 min and then permeabilized with 0.2% triton-X in PBS for 5 min, or fixed and permeabilized at the same time with PEMT buffer (0. 1M PIPES, 1mM MgCl_2_, 1mM EGTA, 0.2% Triton-X, 4% PFA), then blocked with 5% bovine serum albumin in PBS for 30 min and treated with primary and secondary antibodies for 1 hr at room temperature. Primary antibodies were used at the following dilutions: tubulin 1:1000 (DM1A, Sigma-Aldrich), Ensconsin (MAP7, Proteintech, 13446-1-AP) 1:100–200, mCherry 1:500 (Abcam). Secondary anti-rabbit IgG and anti-mouse IgG antibodies (Molecular Probes) tagged with alexa-fluor 488 or 546 were used at 1:500 and DAPI (Invitrogen) at 1:2000. Immuno-stained cells were mounted with FluorSave (Calbiochem) and imaged on a Leica SPE confocal microscope with a 63X lens (NA 1.3) or on a UltraView Vox (Perkin Elmer) spinning disc confocal microscope with 60X (NA 1.4).

### Image Processing and Analysis

Displayed images were processed using ImageJ where necessary, brightness was changed uniformly across the field.

Changes in centrosomal microtubule levels relative to NEP were quantified following way. Median intensity of mEGFP-α-tubulin signal in 15×15 pixel oval around centrosomes and background signal in 3×3 pixel box close to the centrosomes were manually measured in the sum projection images around one centrosome per cell, which did not move a lot in z-direction with ImageJ in the Figure 1D. In the Figure 4B median intensity of mEGFP-α-tubulin signal in 20×20 or 30×30 pixel oval around both centrosomes were measured if centrosomes stayed close enough to each other during the whole course of the movie, otherwise 19×19 pixel oval around one centrosome was used for the measurements. For other data, a custom MATLAB code was used to measure mEGFP-α-tubulin signal at centrosomes semi-automatically. Centrosomes and points for background measurements close to centrosomes were chosen manually and the code calculated the mean of mEGFP-α-tubulin in the cylinder with 15 pixel (for centrosomes) and 5 pixel (for background) diameter and 3 section height. The values displayed in the graphs are background subtracted and normalized by the values at NEP or at −30 min before NEP, as indicated.

Changes in non-centrosomal microtubule levels relative to NEP were quantified following way. Median/variance of mEGFP-α-tubulin signal in 30×30 pixel box at the periphery of a cell and outside a cell (background) was manually measured with ImageJ (Figure S1 A). The values displayed in the graphs are background subtracted and normalized by the values at NEP or at −30 min before NEP, as indicated. In addition, cytoplasmic mEGFP-α-tubulin intensity (value at 6 min after the NEP) was subtracted before normalization (Except in the graphs in Figure S1). In the figure 1 D and Figure 2 B median of mEGFP-α-tubulin was measure only at one location per cell in sum projection images of basal sections. Otherwise, median/variance of mEGFP -α-tubulin was measured at two locations per cell in the single sections.

For nuclear tubulin, median intensity of mEGFP-α-tubulin signal was measured in the box (size indicated in figure legends) in the nucleus and outside the cell in the single sections. The values displayed in the graphs are background subtracted and normalized as indicated in the figures.

To measure tubulin dilution in the cells treated with Nocodazole in non-adherent chambers, median intensity of mEGFP-α-tubulin was measured in 6X6 pixel box at two locations at the periphery of cells and outside the cells (background) in single sections in high resolution movies (60X, spinning disc). In low resolution movies (20X, wide-field microscope) median mEGFP-α-tubulin signal was measured in 4X4 pixel box at four locations at the periphery of the cells and at one location outside the cells (background). The values displayed in the graphs are background subtracted and normalized as indicated in the figures.

To measure changes in MS2-mCherry-NLS concertation the median intensity of MS2-mCherry-NLS signal was measured in 20×20 pixel box at two locations in the nucleus and outside the cell (background) in the single sections. The values displayed in the graphs are background subtracted and normalized as indicated in the figure.

Volume occupied by MS2-mCherry-NLS was measured using a custom MATLAB code, developed to be independent of the overall mCherry intensity. Briefly, a point within the volume occupied by MS2-mCherry-NLS was found using a Difference-of-Gaussians (DoG) filter. This was then expanded to find the boundary of the volume using a gradient watershed algorithm in three dimensions. Cells were tracked from frame to frame using the Munkres algorithm; the minimal movement of cells between frames allowed trivial robust tracking.

To measure tubulin, cytoplasmic mEGFP-α-tubulin intensity upon control, hypo- and hyper-osmotic shock treatments was quantified in following way in 3D: a 30 pixel width line was drawn across the cell in XY and using the FIJI “*KymoResliceWide*” plugin, average intensity values across width was calculated for each section creating kymographies in Z direction for every time point. Then maximum size box according to the cell size was drawn on the kymographs and mean intensities were calculated for every time point.

The maximum diameter of the cells upon control, hypo- and hyper-osmotic shock treatments was calculated by manually identifying contours of the cells every time point in max projection images and measuring corresponding Ferret’s diameter. Spindle mEGFP-α-tubulin intensity and spindle volume were calculating via rendering spindle mEGFP-α-tubulin signal in 3D using Imaris (Bitplane).

Graphs were produced and statistical analysis (as indicated in the figure legends) performed using GraphPad Prism.

### Movie S1. Failure in removal of Ensconsin from microtubules in prophase delays interphase microtubule disassembly and leads to an abnormal looking mitotic spindle

Flat (Rap1*) HeLa cell stably expressing GFP-α-tubulin (green) and non-phosphorylatable Ensconsin microtubule binding domain (A-EMTB-mCherry, red) during mitotic progression. Scale bar represents 10μm.

### Author contributions

N.M. and B.B. wrote the manuscript. N.M. and B.B. designed experiments. N.M. performed and analysed experiments. H.M. helped performing the experiments with non-adhesive chambers. A. C. wrote the Matlab custom code to analyse some data.

## Acknowledgments

N.M thanks SNF (Swiss National Science Foundation), EMBO and Claudio Corrodi for financial support to conduct the project. B.B. thanks Cancer Research UK for funding, and the MRC for core LMCB support. We are grateful to Thomas Surrey, Patrick Meraldi, Ivana Gasic and Nitya Ramkumar for critical reading of the manuscript; to members of B.B. lab for helpful discussions; to J. Pines and S. Wieser for providing wt-EMTB-mCherry plasmid; to E. Paluch for providing LifeAct-GFP; to M. Piel for providing MS2-mCherry-NLS plasmid and hosting N.M in his lab to try the volume measurement method; to Andrew Vaughan for his microscopy expertise.

---

**Figure S1.**
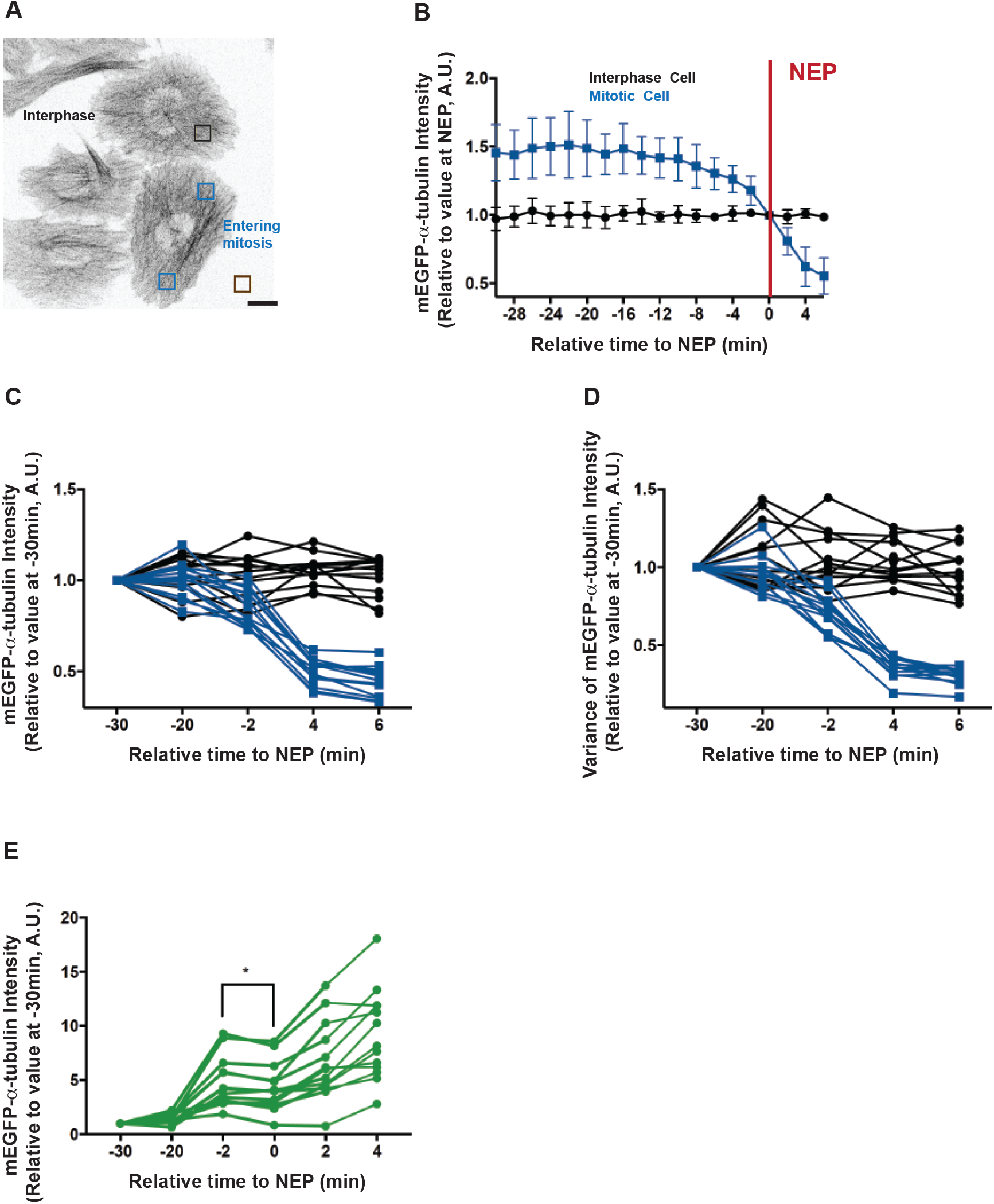
Related to Figure 1. Disassembly of interphase microtubules begins prior to NEP and accelerates at NEP. **A**) Representative confocal image (x-y maximum projection) of H2B-mRFP (not shown) mEGFP-α-tubulin HeLa cells (inverted grayscale) transiently overexpressing Rap1* entering mitosis (the same cell as in Figure 1) and a control cell that remains in interphase during the course of the movie. Inserts show regions that were used for quantifications in B-D. Scale bar represents 10μm. **B**) Changes in non-centrosomal microtubule levels in the cells entering mitosis (blue, the same as in Figure 1D) vs. interphase cells (black, 5 cells from 2 independent experiments). Median intensity of mEGFP-α-tubulin signal was measured as in Figure 1 D. Graph shows means and SD. Changes in non-centrosomal microtubule levels in the cells entering mitosis (blue, the same as in Figure 1 E) vs. interphase cells (black, 13 cells from 4 independent experiments). Median (**C**) and Variance (**D**) of mEGFP -α-tubulin signal was measured as in Figure 1 E. **E**) Changes in centrosomal microtubule levels. Mean of α-tubulin-GFP signal at the centrosomes was measured as in Figure 2 B in the cells entering mitosis at 30, 20, 2 min before NEP and 4, 6 min after NEP (13 cells, 4 independent experiments, the same cells as in Figure 1). Repeated Measures ANOVA with the Greenhouse-Geisser correction, Tukey’s multiple comparisons test with individual variances computed for each comparison. Shown * P=0.039.

**Figure S2.**
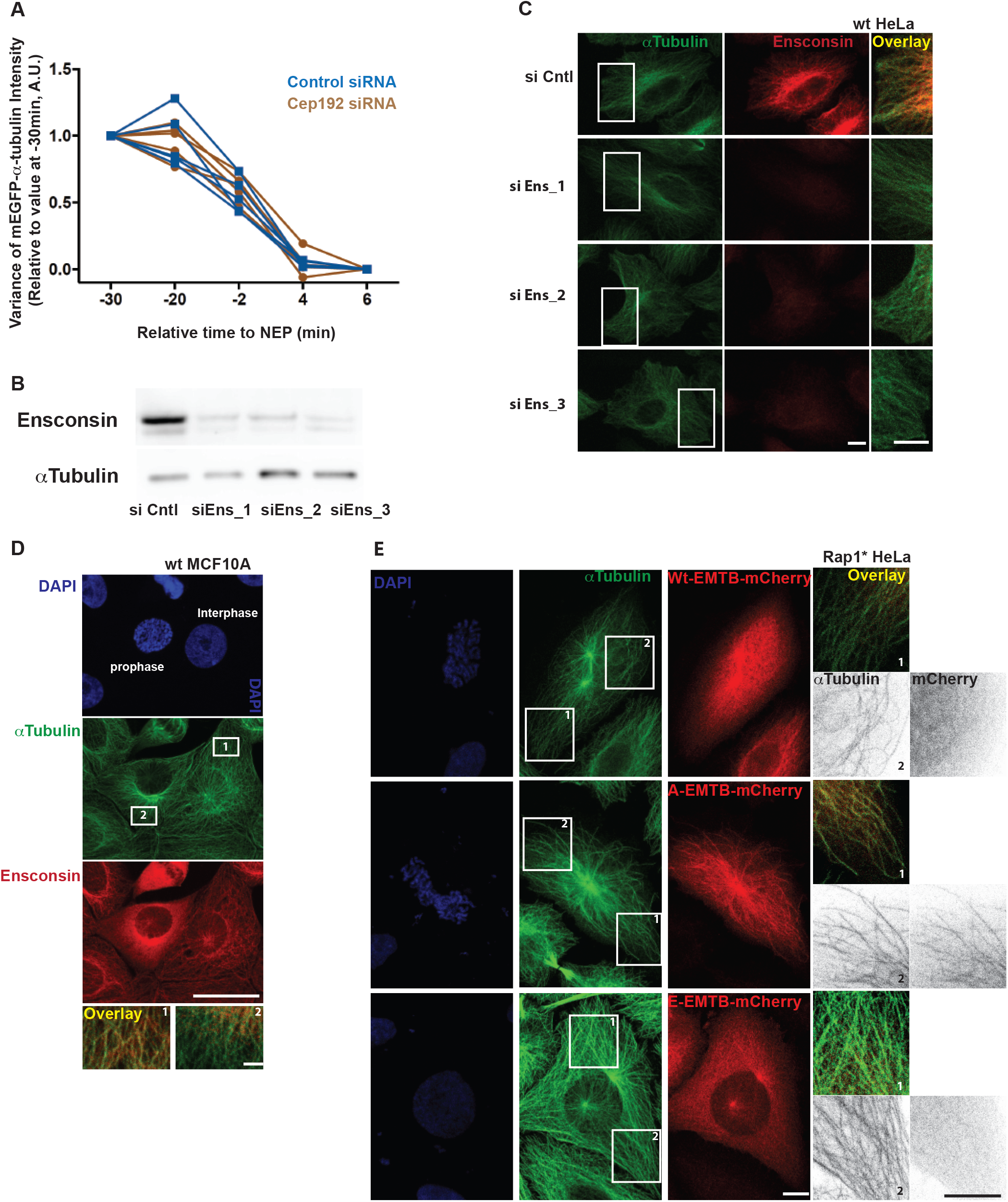
Related to Figure 2. Binding of Ensconsin to microtubules at mitotic entry is regulated by phosphorylation. **A**) Changes in non-centrosomal microtubule levels relative to NEP. Variance of α-tubulin-GFP signal measured as described in Figure S1 D for control siRNA (blue, 5 cells) and Cep192 siRNA cells (brown, 5 cells) 2 experiments as in Figure 2 A-C. **B**) Western blot showing Ensconsin knockdown induced using three different siRNAs targeting Ensconsin. **C**) Representative confocal images (x-y maximum projection) of fixed HeLa cells treated with control siRNA and three siRNAs targeting Ensconsin. Inserts show regions zoomed in overlays (>10 cells per condition, 1 experiment). **D**) Representative confocal images (x-y maximum projection) of fixed MCF10A cells stained to show that Ensconsin is removed from microtubules in prophase compared to interphase. Ensconsin in red, αTubulin in green and DAPI in blue. Inserts show regions zoomed in overlays, in which intensities were adjusted to remove cytoplasmic background signal (6 prophase cells, >20 interphase cells,1 experiment). **E**) Representative confocal images of fixed HeLa cells overexpressing Rap1* in prophase stained to show that the microtubule binding domain of Ensconsin (Wt-EMTB-mCherry) as well as a corresponding phospho-mimetic mutant (E-EMTB-mCherry) are largely cytoplasmic in prophase, whereas the non-phosphorylatable form (A-EMTB-mCherry) localizes to the microtubules. αTubulin in green, Wt-EMTB-mCherry, A-EMTB-mCherry, E-EMTB-mCherry in red. Inserts are zoomed, shown in inverted greyscale or in overlays, where signal intensities were adjusted to remove cytoplasmic background signal (3 cells per condition, 1 experiment). Scale bars represent 10μm.

**Figure S3.**
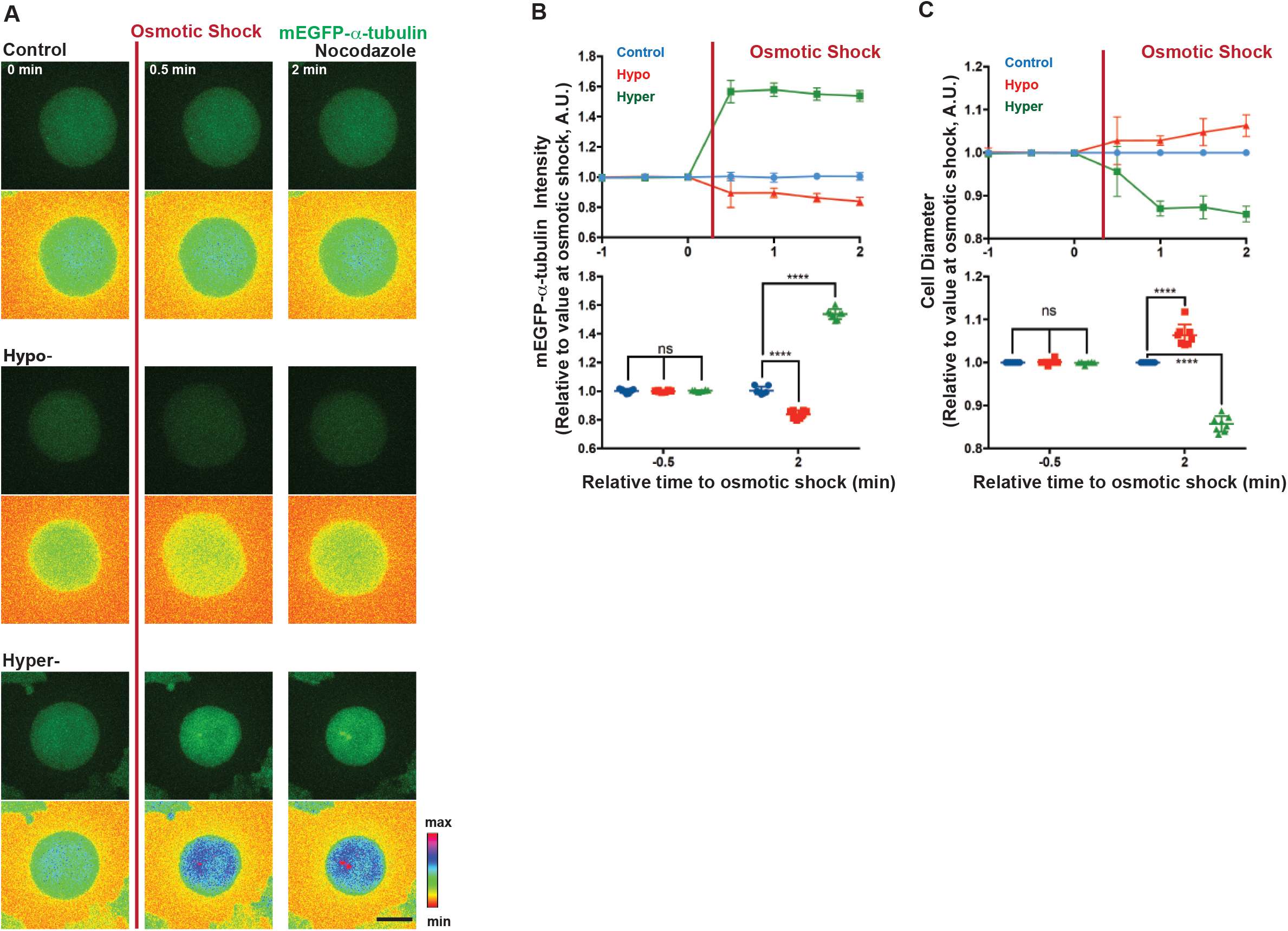
Related to Figure 5. Hypo-osmotic shock can be used to mimic changes in tubulin concentration induced by NEP. **A**) Representative time-lapse confocal images (x-y maximum projection, lower panel: pseudocolor, spectra LUT) of HeLa cells stably expressing H2B-mRFP (was not imaged) and mEGFP-α-tubulin, treated with Nocodazole, to show changes in cell diameter and in mEGFP-α-tubulin intensity before and after hypo- or hyper-osmotic shock treatment relative to control treatment. Scale bar represents 10μm. **B**) Quantifications of changes in mEGFP-α-tubulin intensity induced by osmotic shock relative to control treatment. Mean intensity of mEGFP-α-tubulin signal was measured in cells before and after control (blue, 7 cells), hypo-(red, 8 cells) or hyper- (green, 8 cells) osmotic shock treatments (2 independent experiments) as described in Materials and Methods. **C**) Quantifications of changes in cell diameter (Ferret’s diameter) induced by osmotic shock relative to control treatment. Ferret’s diameter was measured as described in Materials and Methods in cells before and after control (blue, 7 cells), hypo-(red, 8 cells) or hyper-(green, 8 cells) osmotic shock treatments (2 independent experiments). Lower panels show comparison between values at −0.5 min and 2 min relative to osmotic shock treatments. Repeated Measures two-way ANOVA, Dunnett’s multiple comparisons test, **** P=0.0001

